# A metapopulation model for the 2018 Ebola virus disease outbreak in Equateur province in the Democratic Republic of the Congo

**DOI:** 10.1101/465062

**Authors:** Sophie R. Meakin, Mike J. Tildesley, Emma Davis, Matt J. Keeling

## Abstract

Ebola virus disease (EVD) is a viral haemorrhagic fever with high mortality that has caused a number of severe outbreaks in Central and West Africa. Although the majority previous outbreaks have been relatively small, the result of managing outbreaks places huge strains on already limited resources. Mathematical models matched to early case reporting data can be used to identify outbreaks that are at high risk of spreading. Here we consider the EVD outbreak in Equateur Province in the Democratic Republic of the Congo, which was declared on 8 May 2018. We use a simple stochastic metapopulation model to capture the dynamics in the three affected health zones: Bikoro, Iboko and Wangata. We are able to rapidly simulate a large number of realisations and use approximate Bayesian computation, a likelihood-free method, to determine parameters by matching between reported and simulated cases. This method has a number of advantages over more traditional likelihood-based methods as it is less sensitive to errors in the data and is a natural extension to the prediction framework. Using data from 8 to 25 May 2018 we are able to capture the exponential increases in the number of cases in three locations (Bikoro, Iboko and Wangata), although our estimated basic reproductive ratio is higher than for previous outbreaks. Using additional data until 08 July 2018 we are able to detect a decrease in transmission such that the reproductive ratio falls below one. We also estimate the probability of transmission to Kinshasa. We believe this method of fitting models to data offers a generic approach that can deliver rapid results in real time during a range of future outbreaks.

## 1 Introduction

Ebola virus disease (EVD) is a severe viral haemorrhagic fever with a high case fatality ratio ranging from 25% to 90% (World Health Organisation, 2018c). EVD is caused by infection with one of the six known ebolaviruses (family *Filoviridae*), four of which have led to known EVD outbreaks in humans (World Health Organisation, 2018c; Goldstein et al., 2018). Transmission occurs through direct contact with blood or other bodily fluids of symptomatic individuals, or with contaminated materials such as bedding, clothing or needles (World Health Organisation, 2018c). Healthcare workers and caregivers within the community are at high risk of infection (World Health Organisation, 2018c; Evans et al., 2015); transmission can also occur during traditional burial ceremonies that involve direct contact with the body (Hewlett and Amolat, 2003; Victory et al., 2015; Khan et al., 1999).

The 2018 outbreak of EVD in Equateur Province was the twenty-sixth outbreak globally and the ninth within the the Democratic Republic of the Congo (DRC). On 3 May 2018, 21 suspected cases of EVD, including 17 deaths, were identified in the Ikoko-Impenge health area within Bikoro Health Zone in Equateur Province and reported to the DRC Ministry of Health. On 5 May, five active cases in hospitalised patients were identified and samples sent for laboratory testing at the Institute National de Recherche Biomédicale in Kinshasa; of these five active cases, two tested positive for *Zaire ebolavirus*. The outbreak was officially declared by WHO on 8 May 2018. Additional cases were subsequently reported in Iboko and Wangata health zones, the latter of which is located within Mbandaka, a city on the Congo river with a population of approximately 1,2000,000 people (World Health Organisation, 2018d) (Figure 1).

**Figure 1.**
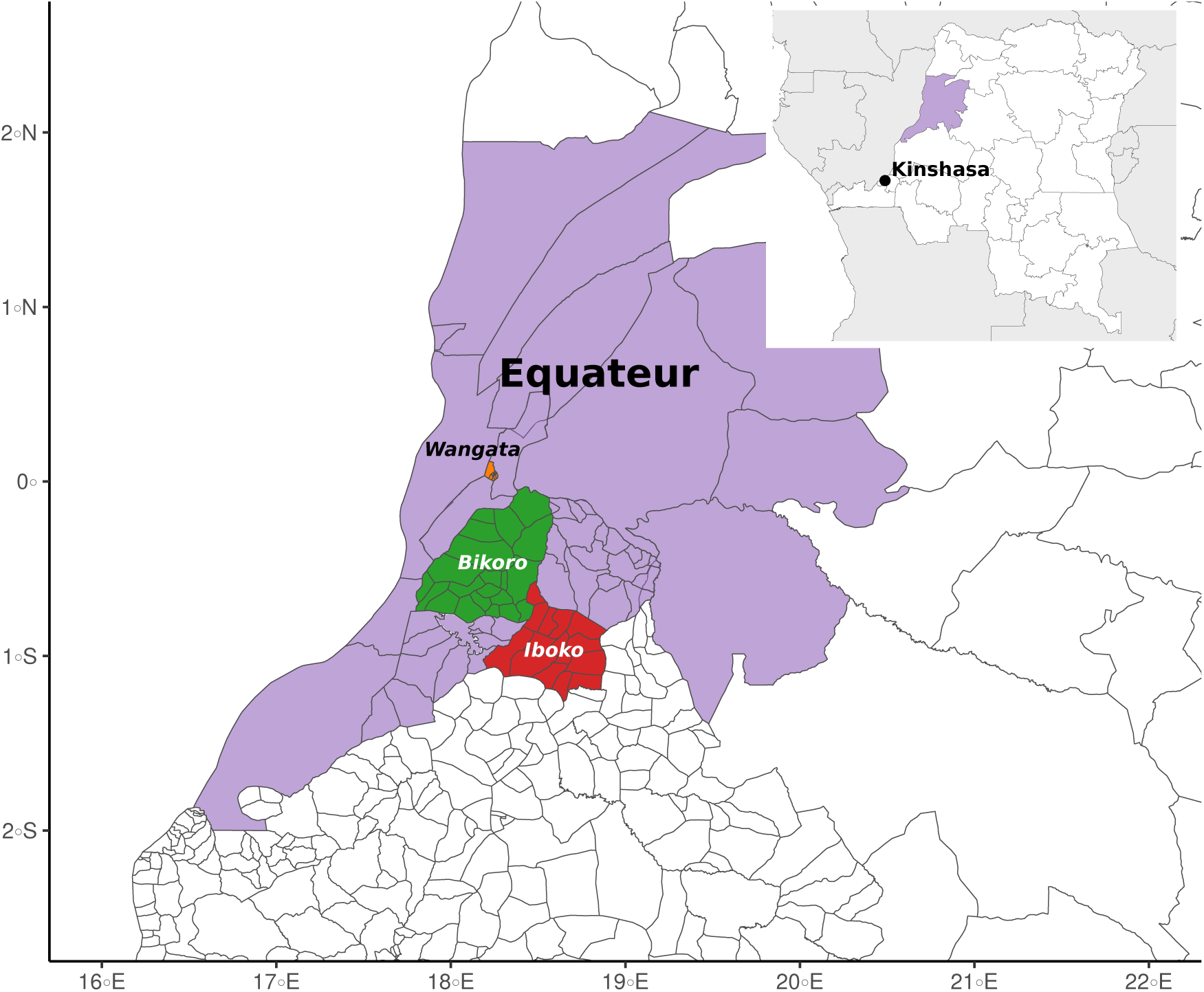
Location of the 2018 EVD outbreak in Equateur Province (purple) in the Democratic Republic of the Congo, highlighting the three affected health zones: Iboko (red), Bikoro (green) and Wangata (yellow). Health zones are further subdivided into health areas. Data from Référentiel Géographique Commun (RGC), DRC Ministry of Health, OpenStreetMap, UCLA, and WHO. Borders do not necessarily reflect official legal boundaries.

There was a rapid local and international response in an effort to prevent further transmission within or out of the DRC. In particular, there was considerable concern over the outbreak reaching the capital city Kinshasa, which has a population of approximately 11,000,000 people and serves international flights to countries in Africa and Europe. Organisations within the affected regions implemented various control measures, including: enhanced community surveillance; the use of rapid diagnostic tests; contact identification and tracing; and safe and dignified burials to reduce possible transmission during funerals. In addition, an experimental EVD vaccine rVSV-ZEBOV was approved by the WHO under compassionate use (World Health Organisation, 2018e), and a ring vaccination strategy began on 21 May 2018. By the end of the outbreak a total of 3,481 individuals had been vaccinated (World Health Organisation, 2018a).

Real-time mathematical modelling can provide important guidance to public health bodies during EVD outbreaks; this information may be particularly beneficial in low-resource settings to target limited resources to regions of greatest need. During the 2014-16 EVD epidemic in West Africa, real-time modelling was used to: forecast case numbers (Rivers et al., 2014; Fisman et al., 2014; Camacho et al., 2015; Merler et al., 2015); predict demand for resources, such as bed requirements (Camacho et al., 2015); assess intervention strategies (Rivers et al., 2014; Ajelli et al., 2015; Merler et al., 2015); and quantify risk of international transmission (Gomes et al., 2014). More recently, Funk et al. (2019) have evaluated the performance of various models used in real-time epidemic forecasting during the West Africa epidemic. An epidemiological study of the 2018 EVD outbreak in Equateur province identified possible routes of exposure and considered the delay between symptom onset, hospitalisation, and sample testing (Barry et al., 2018).

Epidemiological models of EVD often explicitly account for transmission in the community, hospitals and funerals using a six-compartment model introduced by Legrand et al. (2007); this framework has been used, or adapted, by previous real-time EVD modelling studies (Rivers et al., 2014; Gomes et al., 2014; Merler et al., 2015). For the outbreak in Equateur province, previous hospitalisation and funeral attendance were identified as possible routes of exposure for 37% and 60% of total cases, respectively (Barry et al., 2018). Due to the close contact required for onward transmission and the severity of the symptoms, cases of EVD are typically spatially clustered, with occasional long-distance transmission as a result of human movement: for the outbreak in Equateur province, cases in Bikoro and Iboko health zones were restricted to a small number of remote villages (World Health Organisation, 2018a). Agent-based (Merler et al., 2015) and metapopulation modelling (Gomes et al., 2014) have been used to capture spatial dynamics of EVD spread, although parametrisation of agent-based models appear to require highly detailed sociodemographic data and computationally expensive Markov chain Monte Carlo simulations. Metapopulation models typically require fewer resources and so are an excellent framework to use in real-time outbreak modelling.

We adapt the widely-used six-compartment model of EVD dynamics with a non-constant transmission parameter to assess early growth of cases and to identify changes in transmission as the outbreak progresses. We use a metapopulation framework to account for spatial clustering of cases and to allow us to quantify the risk of case importation to Kinshasa.

## 2 Methods

### 2.1 Data

We used reports from the DRC Ministry of Health (DRC Ministère de la Santé, 2018) and WHO (World Health Organisation, 2018b) to produce time series of the number of cases at the health zone level. In both reports, cases are classified as suspected, probable or confirmed according to WHO guidelines (World Health Organisation, 2014), although cases could undergo reclassification at a later date. Due to the uncertainty of the true status of both suspected and probable cases and the possibility of reclassification of these cases, we consider cumulative confirmed cases only. We stress that these data only give the date of laboratory confirmation, not the date of symptom onset or case detection, and therefore these dates are multiple steps removed from the underlying epidemiological dynamics. A time series of the cumulative data for the three health zones is given in Figure 2.

**Figure 2.**
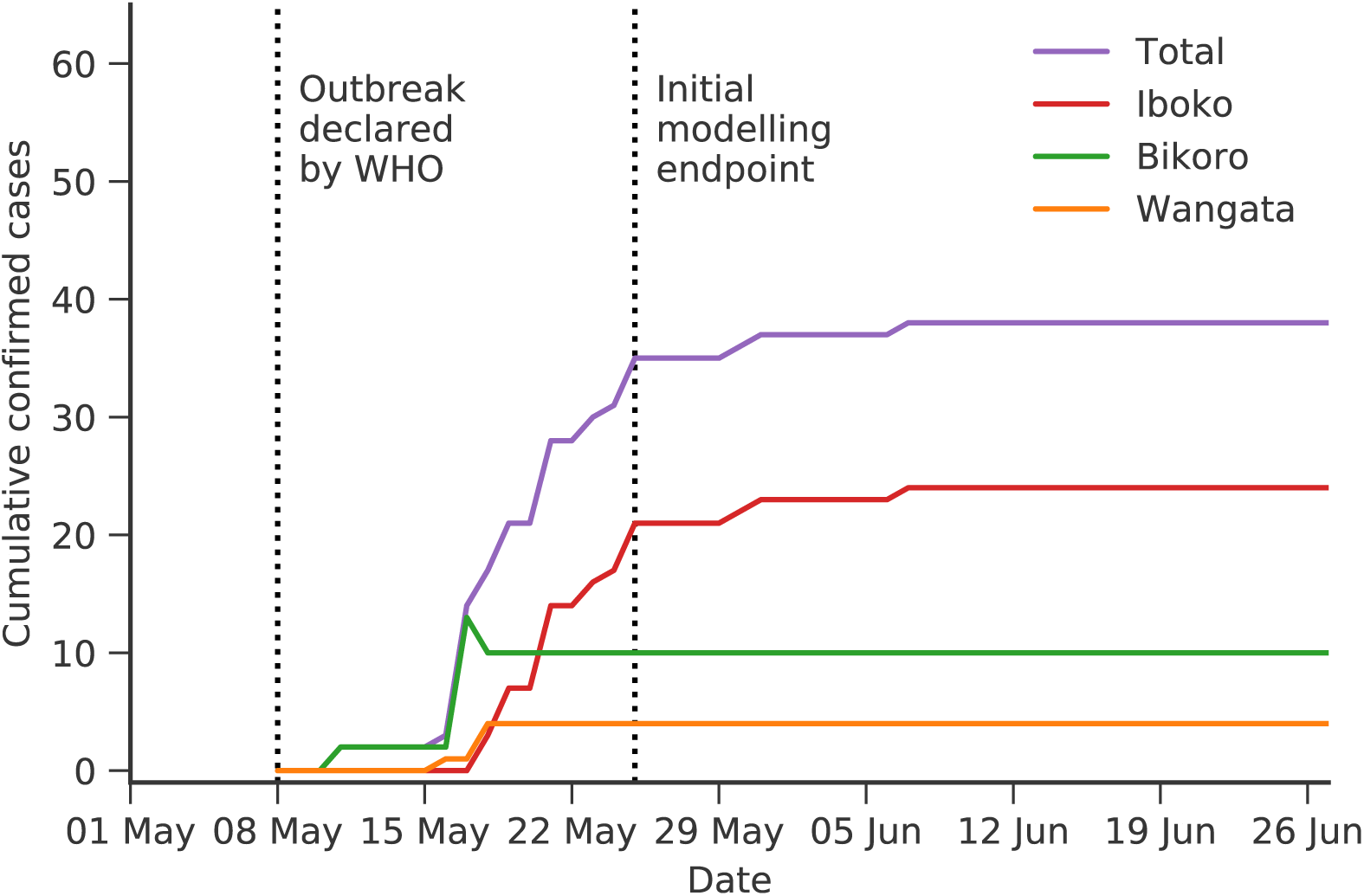
Time series of cumulative confirmed cases in the 2018 EVD outbreak in Equateur Province in the DRC. The outbreak was officially declared by WHO on 8 May 2018. We also indicate the initial modelling endpoint on 25 May 2018.

### 2.2 Model

Our model can be described in two parts: a compartmental model that describes the epidemiological dynamics of EVD within a population (Legrand et al., 2007); and a metapopulation model that describes the spatial dynamics (Keeling and Rohani, 2002).

#### 2.2.1 Epidemiological dynamics

We use a stochastic six-compartment model to describe the epidemiological dynamics of EVD (Legrand et al., 2007), where the six compartments represent: susceptible individuals (*S*), who can be infected after contact with infectious individuals; exposed individuals (*E*), who are infected but not yet infectious to others; infectious individuals (*I*) within the community; hospitalised infectious individuals (*H*); dead individuals (*F*), who are still infectious and may transmit infection during burial; and removed individuals (*R*), who are either recovered or dead and safely buried. Figure 3 shows a schematic representation of the model, while Table 1 summarises the possible events and transition rates and Table 2 summarises the parameter values used in the model. We also introduce an additional scaling parameter 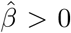 that scales each of the transmission parameters *β*_*I*_, *β*_*H*_ and *β*_*F*_, associated with transmission in the community, hospitals and funerals.

**Table 1.** A summary of possible transitions and their rates for the compartmental model used to describe the epidemiological dynamics of EVD within a population of size *N*. There are ten free parameters in the model and five additional parameters that are calculated from the first ten. See Table 2 for parameter definitions and values.

| Event | Transition | Rate |
| --- | --- | --- |
| Infection | $(S, E) \rightarrow (S - 1, E + 1)$ | $\hat{\beta}S(\beta_I I + \beta_H H + \beta_F F)/N$ |
| Symptom onset | $(E, I) \rightarrow (E - 1, I + 1)$ | $\alpha E$ |
| Hospitalisation | $(I, H) \rightarrow (I - 1, H + 1)$ | $\theta_1 \gamma_{IH} I$ |
| Death (community) | $(I, F) \rightarrow (I - 1, F + 1)$ | $\delta_1 (1 - \theta_1) \gamma_{IF} I$ |
| Recovery (community) | $(I, R) \rightarrow (I - 1, R + 1)$ | $(1 - \delta_1)(1 - \theta_1) \gamma_{IR} I$ |
| Death (hospital) | $(H, F) \rightarrow (H - 1, F + 1)$ | $\delta_2 \gamma_{HF} H$ |
| Recovery (hospital) | $(H, R) \rightarrow (H - 1, R + 1)$ | $(1 - \delta_2) \gamma_{HR} H$ |
| Burial | $(F, R) \rightarrow (F - 1, R + 1)$ | $\gamma_{FR} F$ |

**Table 2.** A summary of the epidemiological parameters used in the compartmental model.

| Parameter |  | Value | Source |
| --- | --- | --- | --- |
| $\beta_I$ | Community transmission rate | 0.588 weeks<sup>-1</sup> | (Legrand et al., 2007) |
| $\beta_H$ | Hospital transmission rate | 0.794 weeks<sup>-1</sup> | (Legrand et al., 2007) |
| $\beta_F$ | Funeral transmission rate | 7.653 weeks<sup>-1</sup> | (Legrand et al., 2007) |
| $\alpha^{-1}$ | Mean duration of incubation period | 7 days | (Bwaka et al. (1999); Ndambi et al. (1999); Dowell et al. (1999)) |
| $\gamma_{IH}^{-1}$ | Mean time from onset to hospitalisation | 5 days | (Khan et al., 1999) |
| $\gamma_{IF}^{-1}$ | Mean time from onset to death | 9.6 days | (Khan et al., 1999) |
| $\gamma_{IR}^{-1}$ | Mean time from onset to end of infectiousness for survivors | 10 days | (Dowell et al. (1999); Rowe et al. (1999)) |
| $\gamma_{FR}^{-1}$ | Mean time from death to traditional burial | 2 days | (Legrand et al., 2007) |
| $\theta$ | Proportion of cases hospitalised | 0.8 | (Khan et al., 1999) |
| $\delta$ | Case fatality ratio (CFR) | 81% | (Khan et al., 1999) |
| $\gamma_{HF}^{-1}$ | Mean time from hospitalisation to death | 4.6 days | Appendix A |
| $\gamma_{HR}^{-1}$ | Mean time from hospitalisation to end of infectiousness for survivors | 5 days | Appendix A |
| $\theta_1$ | Rate of transition from infectious to hospitalised | 0.67 | Appendix A |
| $\delta_1$ | Effective CFR in infected class | 80% | Appendix A |
| $\delta_2$ | Effective CFR in hospitalised class | 80% | Appendix A |

**Figure 3.**
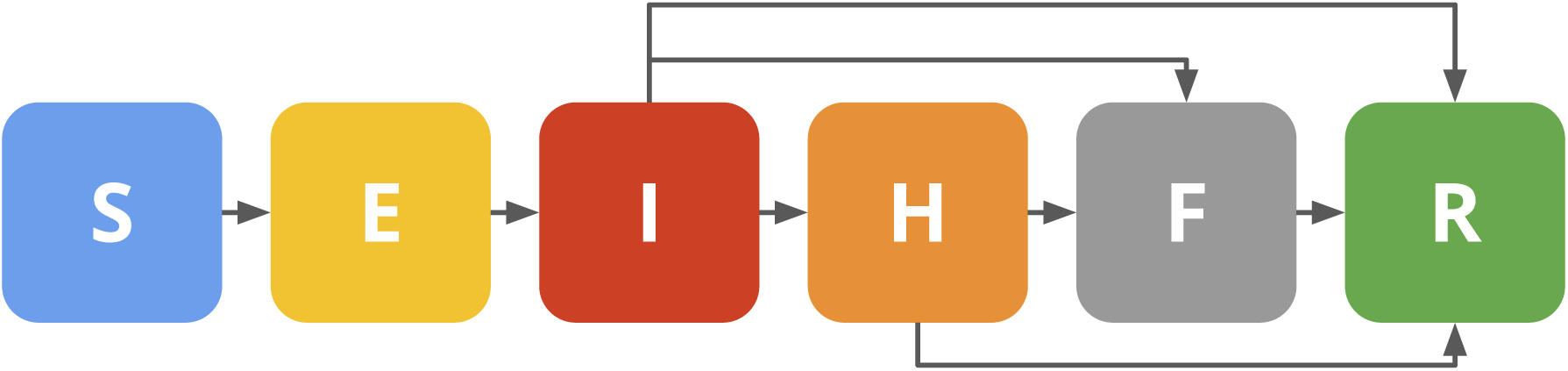
A schematic of the compartmental model used to describe the epidemiological dynamics of EVD within a population, where individuals can be classified as susceptible (*S*), exposed (*E*), infected (*I*), hospitalised (*H*), dead but not yet buried (*F*) or removed (*R*).

##### Model extension

Without change in transmission, our model is likely to predict long-term exponential growth of infection until the susceptible population sizes become depleted; however, sustained exponential growth is rarely observed as intervention measures and individual-level behavioural changes reduce the rate of transmission. We capture possible changes in transmission by including a step-change in 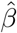 governed by two additional parameters. The transmission scaling, 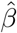 now becomes a function of time such that:

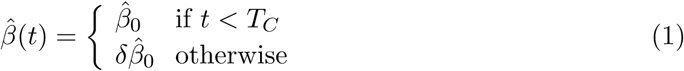

where 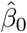 defines the initial transmission scaling, *T*_*C*_ is the time at which control effects begin, and (1 *− δ*) determines the reduction in transmission from all infectious classes.

#### 2.2.2 Spatial dynamics

To describe the spatial dynamics of EVD we use a metapopulation model, whereby the total population is split into *K* interacting sub-populations of sizes *N*_*i*_, *i* = 1*,…, K*. We define *σ_ij_* ∈ [0, 1] to be the proportion of epidemiologically relevant contacts that individuals from population *i* have with individuals in population *j*, which we will simply refer to as the coupling from population *i* to population *j*. We naturally have that 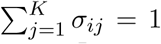, and so the within population coupling can be expressed as *σ*_*ii*_ = 1 − ∑_*j≠i*_ σ_*ij*_. The force of infection in population *i*, the rate at which susceptible individuals become infected, can then be written in terms of the coupling parameters as:

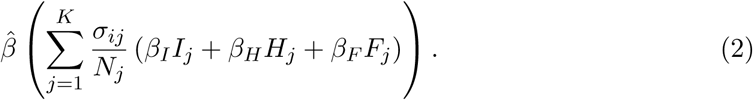

As such the transmission is assumed to be due to the movement of healthy susceptible individuals visiting infected locations, such that the risk to individuals in population *i* is related to the coupling terms *σ*_*ij*_.

We define the coupling according to the generalised gravity model (Xia et al., 2004), where *σ*_*ij*_ depends on the size of populations *i* and *j* and the distance between them:

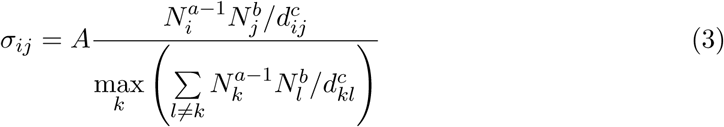

where *N*_*i*_ is the size of population *i*, and *d*_*ij*_ is the straight-line distance between populations *i* and *j*; *a, b, c* and *A* are additional parameters to fit. Population sizes are estimated from census data or, where not available, other online sources (OCHA DR Congo; World Health Organisation, 2018d). Full details of how *σ*_*ij*_ is defined can be found in the Supplementary Information.

### 2.3 Parameter inference

We infer the unknown parameters of the model using approximate Bayesian computation methods (Pritchard et al., 1999). We simulate the outbreak using the tau-leaping algorithm (Gillespie, 2001) and calculate the error between realised (*C*^*sim*^) and observed (*C*^*obs*^) cumulative confirmed cases in each of the sub-populations from day *T*_0_ (to be inferred) for *T*_1_ days; the total error *ε*(*T*_1_) in the *K* sub-populations is calculated as the weighted root mean square error:

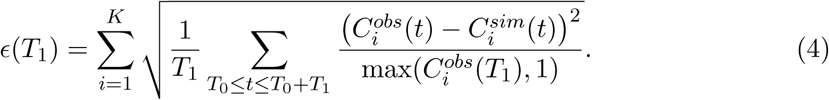

The denominator in this expression is motivated by considering a Poisson distribution. In a Poisson distribution the variance is equal to the mean, therefore we would normalise by dividing through by the observed value at each point; however, given the associated uncertainties in the data, this placed far too much emphasis on correctly matching to the early dynamics when the cases were low. We therefore normalise by the maximum of the observed cases in each location, providing some degree of normalisation between the different sized outbreaks. This approach would fail for Kinshasa where no cases were reported, so we take the normalisation constant to be one.

Parameter inference is performed as a two step process. Parameter values are initially chosen from uniform prior distributions and from 10^7^ parameter sets we retain the 1,000 sets with the lowest error. We then choose parameters based on the current best 1,000 parameter sets: the parameter space to test is normally distributed around each of the best 1,000 parameter sets. In this way, the top 1,000 are updated to reflect newly-tested parameters. The prior distributions used in the first step of the parameter fitting can be found in the Supplementary Information.

### 2.4 Analyses

We first use the model to explore the early growth of cases in the three affected health zones. Using the simple model for *K* = 3 sub-populations (Iboko, Bikoro and Wangata) with data up to 25 May 2018 (equivalent to *T*_1_ = 14 days) we infer the start date *T*_0_, the initial transmission scaling parameter 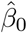 and the between-population coupling.

Next we use the model to identify changes in transmission-as various public-health measures come into effect then we would expect an overall decrease in transmission. Using the breakpoint transmission model for *K* = 3 sub-populations (Iboko, Bikoro and Wangata), we infer the start date *T*_0_, the three transmission parameters (initial transmission scaling 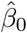, time of change in transmission *T*_*C*_ and the percentage reduction in transmission (1 − *δ*)), and the pairwise between-population coupling; we also estimate the basic reproduction number *R*_0_(*t*) as the transmission changes, given by 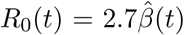, where 2.7 is the estimated *R*_0_ in Legrand et al. (2007). We run this breakpoint transmission model for multiple endpoints, from 25 May 2018 (equivalent to *T*_1_ = 14 days) and in 7 day increments to 6 July 2018 (*T*_1_ = 56 days) to consider how our estimate of *R*_0_ changes as more data is included. We also use these results to estimate the date at which *R*_0_ falls below the threshold for continued transmission (*R*_0_ = 1).

We also use our model with Kinshasa as an additional sub-population to quantify the risk of importation to Kinshasa. Using the breakpoint transmission model for *K* = 4 sub-populations (Iboko, Bikoro, Wangata and Kinshasa), we again infer the transmission parameters and pairwise between-population coupling. We combine these to calculate the cumulative force of infection in each of the sub-populations over time. The force of infection at population *i* at time *t* is given by

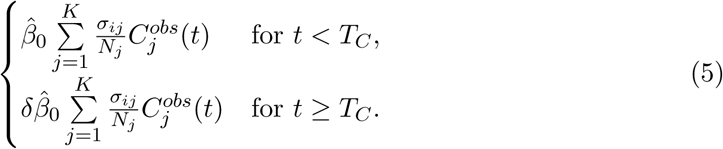

## 3 Results

### 3.1 Early case growth

#### 3.1.1 Initial best-fit parameter estimates

We obtain best-fit estimates of the unknown parameters using data until 25 May 2018 (Figure 4). Firstly, we estimate the start date *T*_0_ and the initial transmission scaling 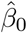. We estimate the start date *T*_0_ to be 25 April 2018 (95% CI [15 April 2018, 03 May 2018]) (Figure 4a). We estimate the initial transmission scaling 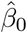 to be 2.7 (95% credible interval (CI) [1.6, 4.5]; that is, the interval that contains 95% of all the parameter values). This gives an initial estimate for the basic reproduction number *R*_0_ of 7.3 (95% CI [4.2, 12.0]) (Figure 4b). Similar parameter values are also obtained for the simple model including Kinshasa as a fourth sub-population.

**Figure 4.**
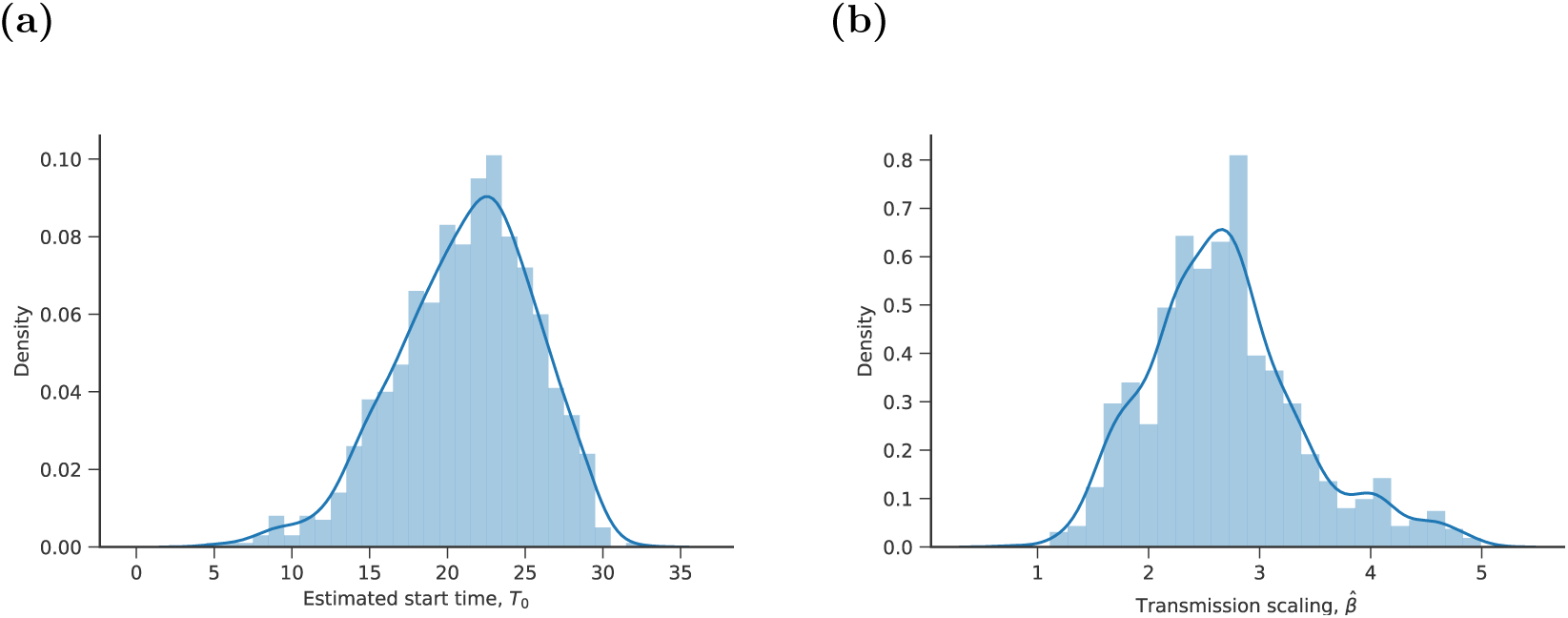
Posterior distributions determined by the 1,000 realisations with the smallest total error fitted to data from 5 April to 25 May 2018 for (a) the start date *T*_0_, (b) the initial transmission scaling parameter 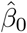.

We recombine the four spatial parameters (*a, b, c* and *A*) as described in Equation (3) to obtain meaningful distributions of the coupling between the sub-populations (Supplementary Information, Figure 1). From these results we observe that the coupling between populations is primarily dominated by distance: the largest coupling is between the closest populations, Bikoro and Iboko. In addition, we find that the coupling is slightly larger towards the bigger population: the coupling from Iboko to Bikoro is larger than from Bikoro to Iboko, since the population size of Bikoro is larger than Iboko.

#### 3.1.2 Best-fit time series

From the model fitting process we also obtain time series fits to the observed cumulative cases and final distribution of cases corresponding to 25 May 2018 (Supplementary Information, Figure 2). We obtain a good qualitative fit to the observed confirmed cases both at the sub-population and metapopulation scale. The individual replicates are tightly clustered and generally envelope the cumulative reported cases. Our model noticeably overestimates the reported number of confirmed cases early in the outbreak; we believe that this overestimate is due to the process by which cases are confirmed. In general, the model estimates improve over time.

### 3.2 Changes in transmission

#### 3.2.1 Best-fit parameter estimates

We obtain best-fit estimates of the unknown parameters as additional confirmed cases are reported (Figure 5). As additional data are used in the model fitting process, we estimate an earlier start date *T*_0_, a lower initial transmission scaling 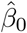 (and hence a lower initial *R*_0_), and a later date at which transmission changes *T*_*C*_. Using data until 6 July 2018 we estimate the start date to be 13 April 2018 (95% CI [05 April 2018, 22 April 2018]) and we estimate the initial transmission scaling to be 1.5 (95% CI [1.1, 1.8]); this gives an initial estimate for *R*_0_ of 3.9 (95% CI [2.9, 4.9]). We estimate the date of change in transmission to be 11 June 2018 (95% CI [02 June 2018, 21 June 2018]), and the percentage reduction in transmission, 1 *− δ*, to be 98.7% (95% CI [92.2%, 100%]) (Supplementary Information, Figure 3). Similar values are also obtained for the breakpoint transmission model including Kinshasa as a fourth sub-population.

**Figure 5.**
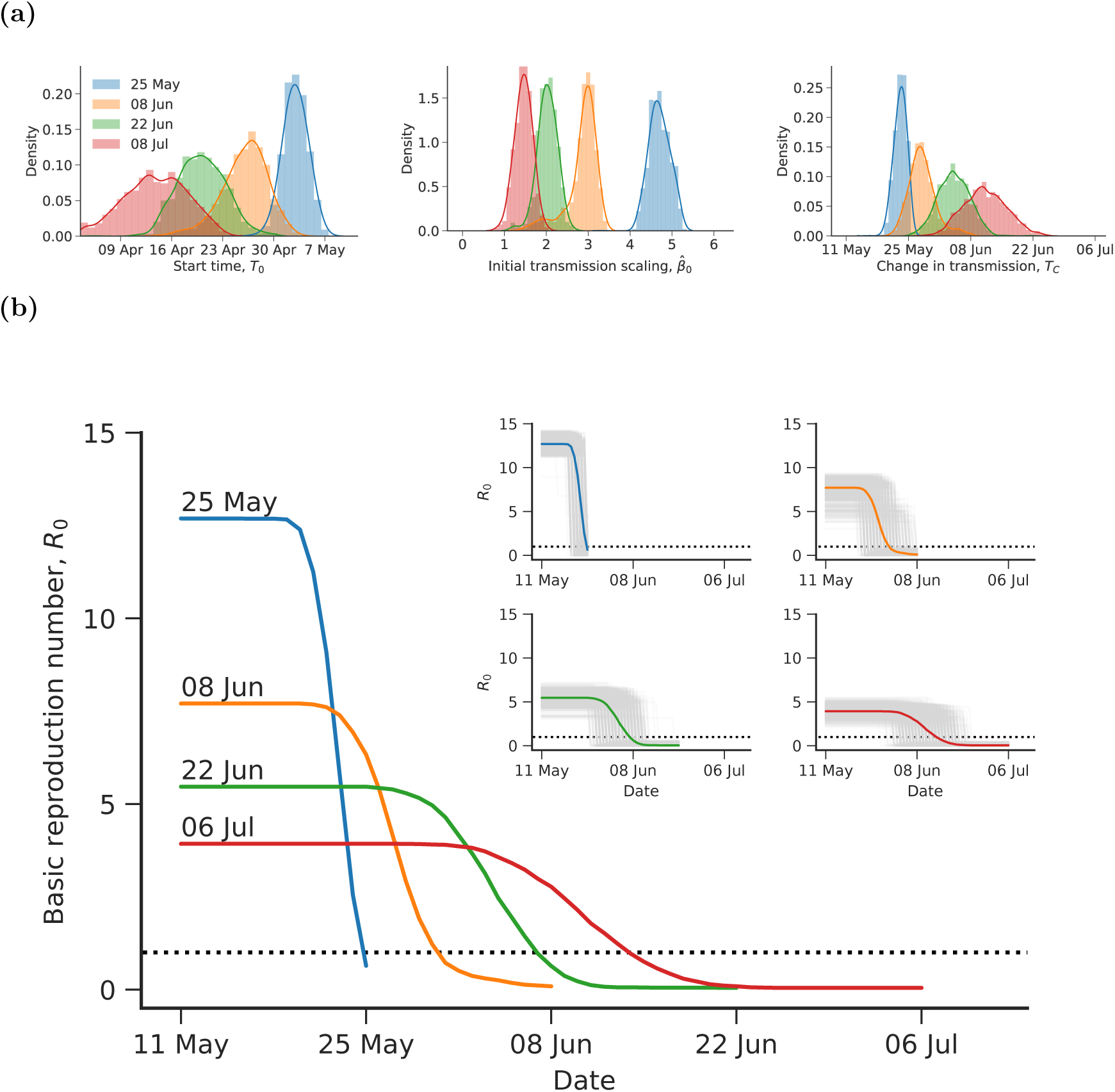
(a) Posterior distributions for the start date *T*_0_, the initial transmission scaling parameter 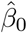 and the date of change in transmission *T*_*C*_. These distributions are determined by the 1,000 best realisations fitted to data up to 25 May, 8 June, 22 June and 6 July.(b) The mean reproductive ratio over time (coloured line) calculated from the 1,000 best realisations fitted to data up to 25 May, 8 June, 22 June and 6 July. The *R*_0_ estimate for the 1,000 best realisations are shown (light grey) for each of the four end dates in the inset figure.

#### 3.2.2 Identifying changes in *R*_0_

We estimate the basic reproductive ratio *R*_0_ as additional confirmed cases are reported and included in the model fitting procedure (Figure 5b). As additional confirmed cases are reported, our initial estimate of *R*_0_ improves. Early in the outbreak (using data until 25 May 2018), we get an initial estimate for *R*_0_ of 12.7 (95% CI [11.5, 13.9])-this is clearly a significant overestimate, but can be attributed to the small amount of data observed, and the process by which cases are confirmed. However, by 6 July 2018 we get an initial estimate for *R*_0_ of 3.9 (95% CI [2.9, 4.9]).

We also estimate the probability that *R*_0_ has fallen below 1, and the time at which this occurs. As additional confirmed cases are reported and included in the model fitting procedure, the probability that *R*_0_ has fallen below 1 increases, and we estimate a later time at which this occurs. Using data until 25 May 2018, the probability that *R*_0_ has fallen below 1 by 25 May 2018 is 0.748 (that is, *R*_0_ has fallen below 1 in 748 out of the 1,000 best realisations). For all successive end dates this probability is 1. Since we are using a simple breakpoint transmission model, the date on which *R*_0_ falls below 1 is the date of change in transmission *T*_*C*_: using data until 6 July 2018, this is 11 June 2018 (95% CI [2 June 2018, 21 June 2018]), approximately 4 weeks after the start of the outbreak.

#### 3.2.3 Best-fit time series

We obtain time series fits to the observed cumulative cases and the final distribution of cases corresponding to 6 July 2018 for the metapopulation as a whole (Figure 6a) and for the four sub-populations separately (Figure 6b). We now more robustly capture the bulk shape of the outbreak including the transition from exponential growth to disease eradication, although we still slightly overestimate the number of confirmed cases during the early stages of the outbreak.

**Figure 6.**
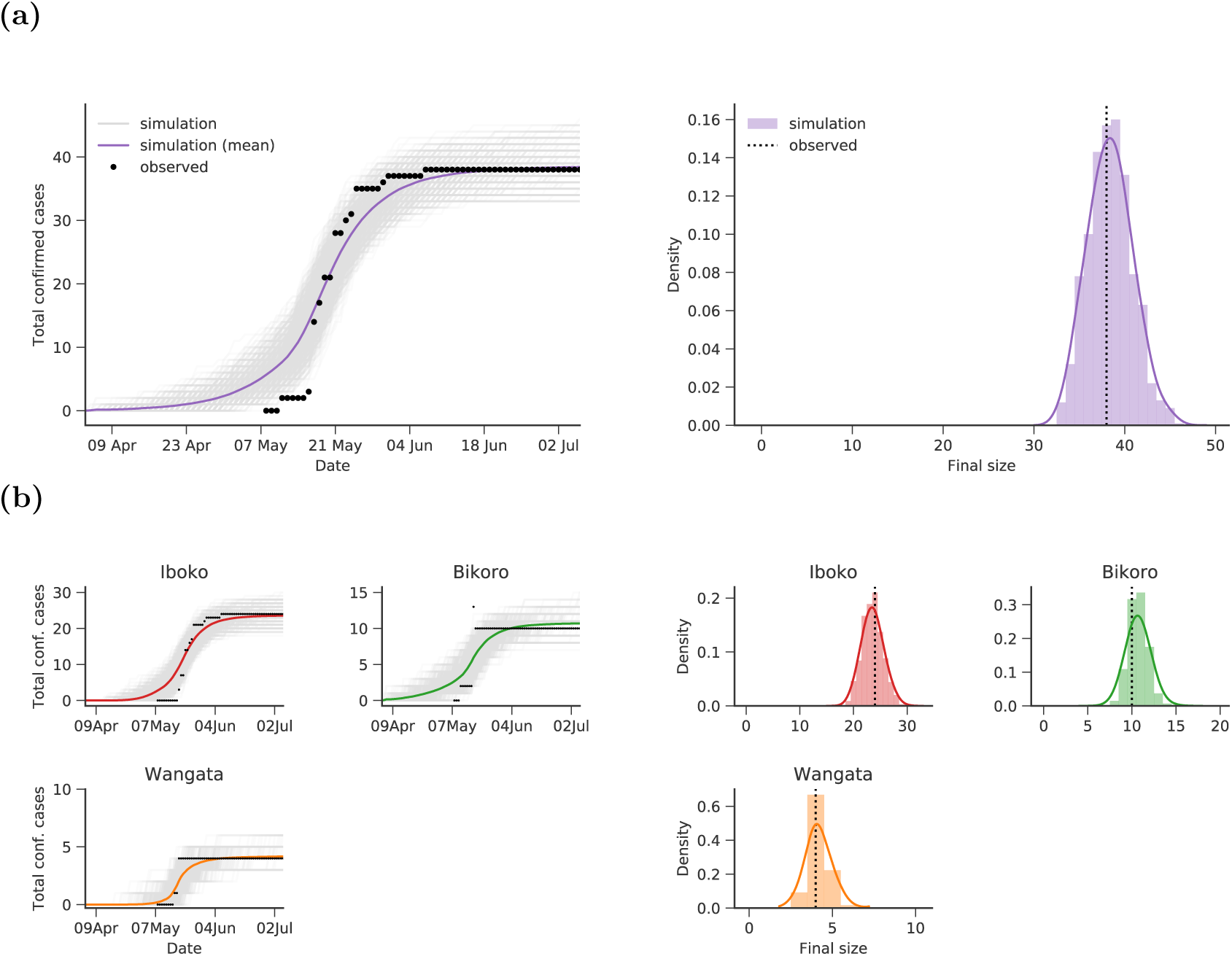
The epidemic curve and final distribution of cases for the 1,000 realisations with the smallest total error for our step-change model up to 6 July 2018. Results are shown for (a) the metapopulation, summed over all sub-populations, and (b) the three sub-populations separately. On the left-hand side we show individual realisations of the outbreak (shown in light grey) plotting the cumulative number of cases moving from the infected to either the hospitalised or funeral class; the mean across all 1,000 realisations is plotted in colour and solid points denote actual cumulative confirmed cases. On the right-hand side we show the distribution of total cases from the 1,000 best realisations together with the reported value (shown as a dashed vertical line).

We compare the final distribution of the realised cumulative confirmed cases to the observed cumulative cases on 6 July 2018 (Figure 6, RHS). We estimate the total number of confirmed cases to be 38 (mean 38.4, 95% CI [33, 44]), matching the 38 observed confirmed cases. At the sub-population level, our mean estimates for the final size are very similar to the observed number of confirmed cases and are more tightly distributed than for the simple model. In Wangata we estimate 4 confirmed cases (mean 4.2, 95% CI [3, 5]), which matches the 4 observed cases; in Bikoro we estimate 11 confirmed cases (mean 10.7, 95% CI [9, 13]) compared to 10 observed cases; and in Iboko we estimate 24 confirmed cases (mean 23.5, 95% CI [20, 27]), matching the 24 observed cases.

### 3.3 Quantifying risk of transmission to Kinshasa

We calculate the cumulative force of infection (Equation 5) in each of the four sub-populations (Figure 7). The force of infection increases as new cases are reported, and decreases around June as transmission is reduced as a result of intervention measures, and plateaus once no new cases are observed. We also observe that the force of infection is highest in Iboko and Bikoro, as we would expect since most of the cases were observed in these two health zones. In comparison, the force of infection in Kinshasa is very small, indicating a very low risk of onward transmission.

**Figure 7.**
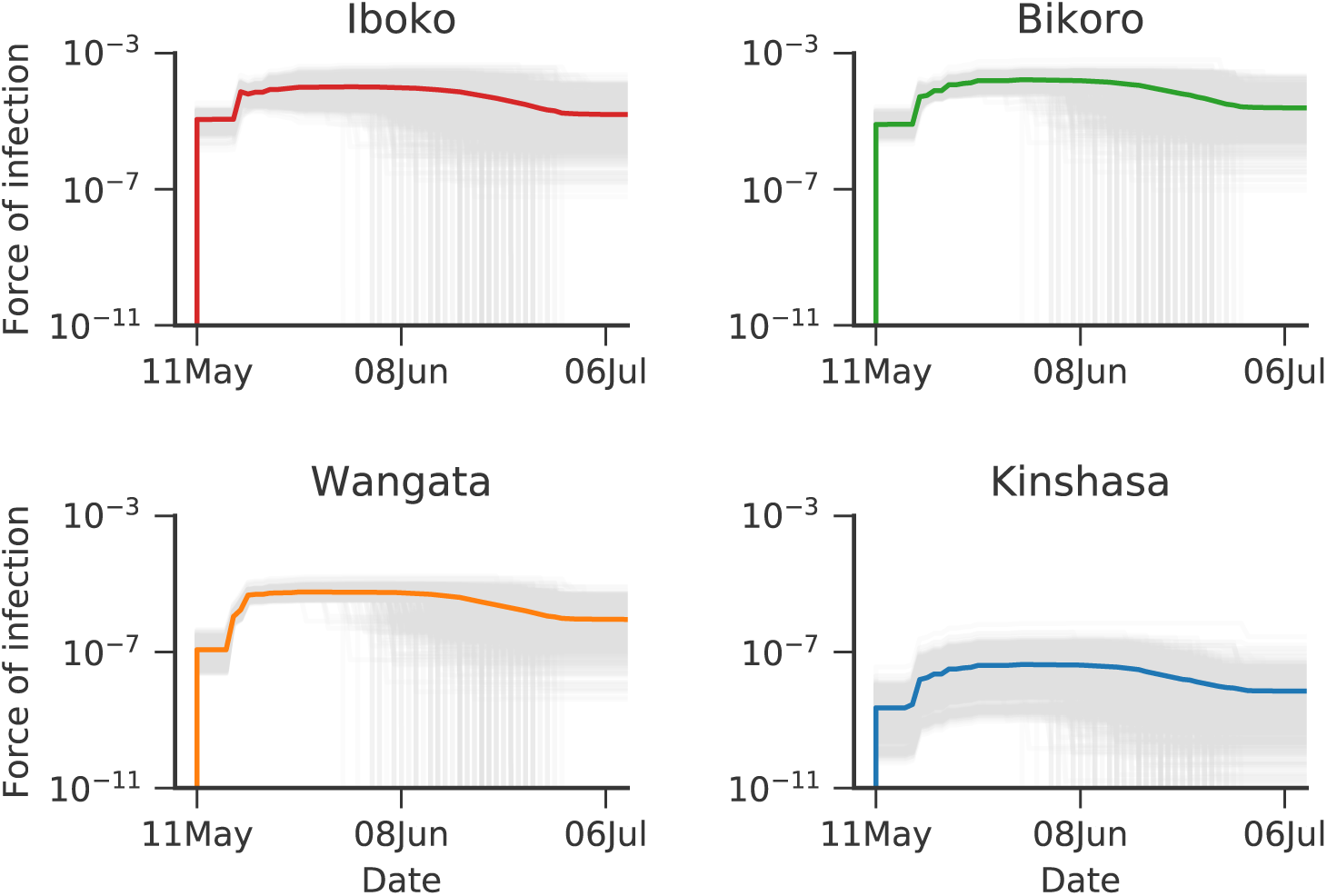
Cumulative force of infection over time in each of the four sub-populations. The mean force of infection taken over all 1,000 realisations is shown as a thick coloured line; individual realisations are shown in light grey.

## 4 Discussion

EVD outbreaks present a significant burden to healthcare resources in countries in Central and West Africa. Real-time mathematical modelling can provide important guidance to public health bodies during EVD outbreaks; this information may be particularly beneficial in low-resource settings to target limited resources to regions of greatest need. We use a six-compartment model of EVD with a non-constant transmission parameter within a metapopulation framework to assess early growth of cases, identify changes in transmission as the outbreak progresses, and to quantify the risk of case importation to Kinshasa.

Our first model is fitted to the start of the outbreak. If we assume constant transmission then we would predict long-term exponential growth, which was not observed; however our second model is able to capture the exponential growth phase and disease eradication. We estimate a 98.7% reduction in transmission and that the basic reproductive ratio *R*_0_ drops below one on 11 June 2018, indicating that the control measures are having their desired impact. The magnitude of the change is in qualitative agreement with WHO reports detailing the extensive local and international response to the outbreak.

Our model combines a well-established compartmental model for EVD (Legrand et al., 2007) with a spatial metapopulation structure. Metapopulation modelling has previously been used to assess the risk of international spread of EVD during the 2014 outbreak in West Africa (Gomes et al., 2014); however, this approach relies upon the Global Epidemic and Mobility Model (Balcan et al., 2010) and hence cannot be easily adapted for use at smaller scales. Given the temporal disconnect between the epidemiologically important infection times and the observed confirmation times, we adopt a likelihood-free approach where repeated stochastic simulations are used to minimise the error between model and data. Our framework has several practical advantages: it can be readily used with any form of stochastic model, and for small outbreak sizes many realisations can be generated quickly, such that many millions of simulations can be performed and analysed.

Although our model is able to capture the dynamics of the outbreak in Equateur Province, the inferred parameter values for 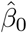, and thus our estimate for *R*_0_, are somewhat surprising. For the simple model without change in transmission for the first two weeks of data we get an estimate for *R*_0_ of 7.3 (95% CI [4.2, 12.0]); for the step-change model using all the data, our new estimate of *R*_0_ is 3.9 (95% CI [2.9, 4.9]). Both values are appreciably larger than other estimates of 1.03 (Barry et al., 2018), and larger than estimates for recent outbreaks: 1.38-3.65 for DRC, 1995; 1.34-2.7 for Uganda, 2000-1 (Camacho et al., 2014); 1.51-2.53 for West Africa, 2013-15 (Van Kerkhove et al., 2015; Althaus, 2014). We believe that this overestimate of *R*_0_ is a result of the process by which cases are confirmed. Due to the delay between symptom onset and laboratory confirmation, confirmed cases are both spatially and temporally clustered, particularly early on in the outbreak. This delay is shown to be longer at the beginning of the outbreak when surveillance is low (Barry et al., 2018). The effect of the delay between symptom onset and laboratory confirmation can be seen as large jumps in the number of confirmed cases in each of Wangata, Bikoro and Iboko. It is unlikely, however, that all cases confirmed on the same day share the same data of symptom onset, which is clear if we compare our data to Barry et al. (2018). This temporal clustering of confirmed cases distorts the data, and is likely the main factor that leads to our overestimate of *R*_0_ compared to previous outbreaks. In principle, we could formulate a model that could mimic the temporal aggregation of cases, but this would generate an additional parameter that would need to be estimated and would place an extra layer of filtering between the epidemiology and results. Alternatively, the overestimate of *R*_0_ could be addressed by modifying and refitting our model to data on the timing of symptom onset, if available.

Our modelling is motivated by the need for real-time analysis of EVD outbreaks and interventions in a spatial setting. Our analysis is constrained by the quality and detail of the limited data publicly available during the outbreak in Equateur Province. In an attempt to minimise uncertainty around the true status of probable and confirmed cases we have restricted our analysis to confirmed cases only. However, even when we only consider confirmed cases the data we use contains at least one error: the number of confirmed cases in Bikoro drops from 13 to 10 on 17 May 2018. Due to the relatively small number of cases and the limited amount of data publicly available, we only infer parameters associated with the spatial component of the model and a single epidemiological parameter that scales the transmission rate; other epidemiological parameters are taken from the literature (Legrand et al., 2007). We believe that inference of all the epidemiological parameters of our model is not possible with the type of data that is publicly available; instead we would require information on the history and treatment of cases at the individual level.

Our quantitative results are also limited by assumptions and approximations made during the modelling process. To define the coupling we use the pairwise straight-line distances between populations; however, straight-line distances are likely a poor proxy for ease of travel between populations: in some remote areas (such as parts of Bikoro and Iboko health zones), it may take a significant amount of time to travel over short distances due to poor road infrastructure. In addition, it is not clear that each of the four sub-populations is acting as a homogeneously mixing population, and hence additional spatial structure may be acting on the dynamics, although without more detailed reporting this is impossible to assess. Modelling at a finer spatial scale would also require additional information on population structure and would increase the computational power and time required for simulation and analysis, which is at odds with our aim to generate results in real-time in low-resource settings.

Our analysis has demonstrated that practically useful mathematical models can be matched to publicly available data early in an outbreak, especially if previous analysis has helped to set the time-course of disease progression. The likelihood-free method we have adopted is highly convenient, allowing us to quickly and easily perform matches between a rapid stochastic simulation model and available data. As such the modelling framework that we have described offers a template for early model inference to other outbreaks. In particular, the framework can easily be modified to accommodate different compartmental models, spatial scales, or data sources. Using this framework, and with very limited publicly-available data, we have been able to attribute a very low risk for the infection reaching Kinshasa which would exacerbate wider dissemination; we have also been able to rapidly identify changes in the transmission rate due to public-health interventions and predict that these interventions are sufficient to curtail the spread of infection. Obviously, as more data become available, especially individual-level data on cases, there is a desire to develop bespoke models fitted to the details of the ensuing outbreak; however, rapid early predictions before too much infection has arisen, such as outlined here, have the potential to generate a substantial impact on public-health decision-making.

## Authors’ contributions

S.R.M, M.J.T and M.J.K. developed the initial concepts; E.D maintained the resarch data; S.R.M and M.J.K performed the computational analysis. All authors played a role in the methodology of the of the research, and in writing and editing the manuscript.

## References

M. Ajelli, S. Parlamento, D. Bome, A. Kebbi, A. Atzori, C. Frasson, G. Putoto, D. Carraro, and S. Merler. The 2014 Ebola virus disease outbreak in Pujehun, Sierra Leone: Epidemiology and impact of interventions. BMC Medicine, 13(1):EE, 2015. ISSN 17417015. doi: 10.1186/s12916-015-0524-z.

C. L. Althaus. Estimating the reproduction number of Ebola virus (EBOV) during the 2014 outbreak in West Africa. PLoS Currents, pages 1–9, 2014. ISSN 2157-3999. doi: 10.1371/currents.outbreaks.91afb5e0f279e7f29e7056095255b288.

D. Balcan, B. GonÇalves, H. Hu, J. J. Ramasco, V. Colizza, and A. Vespignani. Modeling the spatial spread of infectious diseases: The global epidemic and mobility computational model. Journal of Computational Science, 1(3):132–145, 2010. ISSN 18777503. doi: 10.1016/j.jocs.2010.07.002.

A. Barry, S. Ahuka-Mundeke, Y. Ali Ahmed, Y. Allarangar, J. Anoko, B. Nicholas Archer, A. Aruna Abedi, J. Bagaria, M. Roseline Darnycka Belizaire, S. Bhatia, T. Bokenge, E. Bruni, A. Cori, E. Dabire, A. Mouctar Diallo, B. Diallo, C. Ann Donnelly, I. Dorigatti, T. Choden Dorji, A. Rocio Escobar Corado Waeber, I. Socé Fall, N. M. Ferguson, R. Gareth FitzJohn, G. Leon Folefack Tengomo, P. Bernard Henri Formenty, A. Forna, A. Fortin, T. Garske, K. A. Gaythorpe, C. Gurry, E. Hamblion, M. Harouna Djingarey, C. Haskew, S. Alexandre Louis Hugonnet, N. Imai, B. Impouma, G. Kabongo, O. Ilunga Kalenga, E. Kibangou, T. Min-Hyung Lee, C. Okot Lukoya, O. Ly, S. Makiala-Mandanda, A. Mamba, P. Mbala-Kingebeni, F. Fortune Roland Mboussou, T. Mlanda, V. Mondonge Makuma, O. Morgan, A. Mujinga Mulumba, P. Mukadi Kakoni, D. Mukadi-Bamuleka, J.-J. Muyembe, N. Tambwe Bathé, P. Ndumbi Ngamala, R. Ngom, G. Ngoy, P. Nouvellet, J. Nsio, K. Babila Ousman, E. Peron, J. Aaron Polonsky, M. J. Ryan, A. Touré, R. Towner, G. Tshapenda, R. Van De Weerdt, M. Van Kerkhove, A. Wendland, D. Konan Michel Yao, Z. Yoti, E. Yuma, G. Kalambayi Kabamba, J. de Dieu Lukwesa Mwati, G. Mbuy, L. Lubula, A. Mutombo, O. Mavila, Y. Lay, E. Kitenge, and T. Ebola Outbreak Epidemiology Team. Outbreak of Ebola virus disease in the Democratic Republic of the Congo, AprilMay, 2018: an epidemiological study. The Lancet, 392:213–221, 2018. ISSN 01406736. doi: 10.1016/S0140-6736(18)31387-4.

M. A. Bwaka, M.-J. Bonnet, P. Calain, R. Colebunders, A. De Roo, Y. Guimard, K. R. Katwiki, K. Kibadi, M. A. Kipasa, K. J. Kuvula, B. B. Mapanda, M. Massamba, K. D. Mupapa, J. Muyembe Tamfum, E. Ndaberey, C. J. Peters, P. E. Rollin, and E. Van den Enden. Ebola Hemorrhagic Fever in Kikwit, Democratic Republic of the Congo: Clinical Observations in 103 Patients. The Journal of Infectious Diseases, 179(s1):S1–S7, 1999. ISSN 0022-1899. doi: 10.1086/514308.

A. Camacho, A. Kucharski, S. Funk, J. Breman, P. Piot, and W. Edmunds. Potential for large outbreaks of Ebola virus disease. Epidemics, 9:70–78, dec 2014. doi: 10.1016/j.epidem.2014.09.003.

A. Camacho, A. Kucharski, Y. Aki-Sawyerr, M. A. White, S. Flasche, M. Baguelin, T. Pollington, J. R. Carney, R. Glover, E. Smout, A. Tiffany, W. J. Edmunds, and S. Funk. Temporal changes in ebola transmission in Sierra Leone and implications for control requirements: a real-time modelling study. PLoS Currents, 7(OUTBREAKS):1–18, 2015. ISSN 21573999. doi: 10.1371/currents.outbreaks. 406ae55e83ec0b5193e3085.

S. F. Dowell, R. Mukunu, T. G. Ksiazek, A. S. Khan, P. E. Rollin, and C. J. Peters. Transmission of Ebola Hemorrhagic Fever: A Study of Risk Factors in Family Members, Kikwit, Democratic Republic of the Congo, 1995. The Journal of Infectious Diseases, 179(s1):S87–S91, 1999. ISSN 0022-1899. doi: 10.1086/514284.

DRC Ministère de la Santé. Democratic Republic of the Congo Ministère de la Santé mailing list, 2018. URL https://us13.campaign-archive.com/home/?u=89e5755d2cca4840b1af93176{\&}id=aedd23c530.

D. K. Evans, M. Goldstein, and A. Popova. Health-care worker mortality and the legacy of the Ebola epidemic, 2015. ISSN 2214109X.

D. Fisman, E. Khoo, and A. Tuite. Early Epidemic Dynamics of the West African 2014 Ebola Outbreak: Estimates Derived with a Simple Two-Parameter Model. PLoS Currents, pages 1–14, 2014. ISSN 2157-3999. doi: 10.1371/currents.outbreaks.89c0d3783f36958d96ebbae97348d571.

S. Funk, A. Camacho, A. J. Kucharski, R. Lowe, R. M. Eggo, and W. J. Edmunds. Assessing the performance of real-time epidemic forecasts: A case study of the 2013-16 Ebola epidemic. PLoS Computational Biology, 15(2), 2019. ISSN 1553-7358. doi: 10.1101/177451.

D. T. Gillespie. Approximate accelerated stochastic simulation of chemically reacting systems. Journal of Chemical Physics, 115(4):1716–1733, 2001. ISSN 00219606. doi: 10.1063/1.1378322.

T. Goldstein, S. J. Anthony, A. Gbakima, B. H. Bird, J. Bangura, A. Tremeau-Bravard, M. N. Belaganahalli, H. L. Wells, J. K. Dhanota, E. Liang, et al. The discovery of bombali virus adds further support for bats as hosts of ebolaviruses. Nature microbiology, 3(10):1084, 2018.

M. F. C. Gomes, A. Pastore Piontti, L. Rossi, D. Chao, I. Longini, M. Elizabeth Halloran, A. Vespignani, G. Mfc, and P. A. Piontti. Assessing the international spreading risk associated with the 2014 West African Ebola outbreak. PLOS Currents, 6, 2014. doi: 10.1371/currents.outbreaks.cd818f63d40e24aef769dda7df9e0da5.

B. S. Hewlett and R. P. Amolat. Cultural contexts of Ebola in Northern Uganda. Emerging Infectious Diseases, 9(10):1242–1248, 2003. ISSN 10806040. doi: 10.3201/eid0910.020493.

M. J. Keeling and P. Rohani. Estimating spatial coupling in epidemiological systems: a mechanistic approach. Ecology Letters, 5(1):20–29, 2002. ISSN 1461023X. doi: 10.1046/j.1461-0248.2002.00268.x.

A. S. Khan, F. K. Tshioko, D. L. Heymann, B. Le Guenno, P. Nabeth, B. Kerstiëns, Y. Fleerackers, P. H. Kilmarx, G. R. Rodier, O. Nkuku, P. E. Rollin, A. Sanchez, S. R. Zaki, R. Swanepoel, O. Tomori, S. T. Nichol, C. J. Peters, T. G. Ksiazek, and E. A. Poe. The Reemergence of Ebola Hemorrhagic Fever, Democratic Republic of the Congo, 1995. The Journal of Infectious Diseases, page 1842, 1999.

J. Legrand, R. F. Grais, P. Y. Boelle, A. J. Valleron, and A. Flahault. Understanding the dynamics of Ebola epidemics. Epidemiology and Infection, 135(4):610–621, 2007. doi: 10.1017/S0950268806007217.

S. Merler, M. Ajelli, L. Fumanelli, M. F. Gomes, A. P. y. Piontti, L. Rossi, D. L. Chao, I. M. Longini, M. E. Halloran, and A. Vespignani. Spatiotemporal spread of the 2014 outbreak of Ebola virus disease in Liberia and the effectiveness of non-pharmaceutical interventions: A computational modelling analysis. The Lancet Infectious Diseases, 15(2):204–211, 2015. ISSN 14744457. doi: 10.1016/S1473-3099(14)71074-6.

R. Ndambi, P. Akamituna, M.-J. Bonnet, A. M. Tukadila, J. MuyembeTamfum, and R. Colebunders. Epidemiologic and Clinical Aspects of the Ebola Virus Epidemic in Mosango, Democratic Republic of the Congo, 1995. The Journal of Infectious Diseases, 179(s1):S8–S10, 1999. ISSN 0022-1899. doi: 10.1086/514297.

OCHA DR Congo. DR Congo-Health Zones. URL https://data.humdata.org/dataset/dr-congo-health-0.

J. K. Pritchard, M. T. Seielstad, A. Perez-Lezaun, and M. W. Feldman. Population Growth of Human Y Chromosomes: A Study of Y Chromosome Microsatellites. Molecular Biology and Evolution, 16(12): 1791–1798, 1999. ISSN 0737-4038.

C. Rivers, E. Lofgren, M. Marathe, S. Eubank, and B. Lewis. Modeling the Impact of Interventions on an Epidemic of Ebola in Sierra Leone and Liberia. PLoS Currents, 1, 2014. ISSN 2157-3999. doi: 10.1371/currents.outbreaks.fd38dd85078565450b0be3fcd78f5ccf.

A. K. Rowe, J. Bertolli, A. S. Khan, R. Mukunu, J. J. MuyembeTamfum, D. Bressler, A. J. Williams, C. J. Peters, L. Rodriguez, H. Feldmann, S. T. Nichol, P. E. Rollin, and T. G. Ksiazek. Clinical, Virologic, and Immunologic FollowUp of Convalescent Ebola Hemorrhagic Fever Patients and Their Household Contacts, Kikwit, Democratic Republic of the Congo. The Journal of Infectious Diseases, 179(s1): S28–S35, 1999. ISSN 0022-1899. doi: 10.1086/514318.

M. D. Van Kerkhove, A. I. Bento, H. L. Mills, N. M. Ferguson, and C. A. Donnelly. A review of epidemiological parameters from Ebola outbreaks to inform early public health decision-making. Scientific Data, 2:1–10, 2015. ISSN 20524463. doi: 10.1038/sdata.2015.19.

K. R. Victory, F. Coronado, S. O. Ifono, T. Soropogui, B. A. Dahl, C. Centers for Disease, and Prevention. Ebola transmission linked to a single traditional funeral ceremony - Kissidougou, Guinea, December 2014-January 2015. MMWR Morbidity and Mortality Weekly Report, 64(14):386–388, 2015. ISSN 01492195. doi: 10.15585/mmwr.mm6438a2.

World Health Organisation. Case definition recommendations for Ebola or Marburg virus diseases. Technical report, 2014.

World Health Organisation. Ebola virus disease. Democratic Republic of Congo. External Situation Report 17. Technical report, 2018a.

World Health Organisation. Ebola virus disease. Ebola situation reports: Democratic Republic of the Congo., 2018b.

World Health Organisation. Ebola virus disease fact sheet, 2018c. URL http://www.who.int/en/news-room/fact-sheets/detail/ebola-virus-disease.

World Health Organisation. Emergencies preparedness, response. Ebola virus disease-Democratic Republic of the Congo. Disease outbreak news., 2018d. URL http://www.who.int/csr/don/14-may-2018-ebola-drc/en/.

World Health Organisation. Ebola virus disease. Frequently Asked Questions, 2018e. URL https://www.who.int/ebola/drc-2018/faq-vaccine/en/.

Y. Xia, O. N. Bjørnstad, and B. T. Grenfell. Measles metapopulation dynamics: a gravity model for epidemiological coupling and dynamics. The American Naturalist, 164(2):267–281, 2004. doi: 10.1086/ 422341.

