## Supplementary Information for "A metapopulation model for the 2018 Ebola virus disease outbreak in Equateur province in the Democratic Republic of the Congo"

#### 1 Formulation of parameters in epidemic model

There are ten free parameters in the model and five additional parameters that are calculated from the first ten. The model parameters that can be derived from the free parameters are: $\gamma_{HF}^{-1}$, the mean time from hospitalisation to death; $\gamma_{HR}^{-1}$, the mean time from hospitalisation to end of infectious period for survivors; $\theta_1$, the rate of transition from infectious to hospitalised, which is calculated from $\theta$ such that $\theta\%$ of infectious cases are hospitalised; $\delta_1$ and $\delta_2$, the effective case fatality ratios in the infected and hospitalised class, respectively, both which are calculated from $\delta$ such that the case fatality ratio is $\delta$. These five parameters are calculated as follows:

$$\gamma_{HF}^{-1} = \gamma_{IF}^{-1} - \gamma_{IH}^{-1} \tag{1}$$
$$\gamma_{HR}^{-1} = \gamma_{IR}^{-1} - \gamma_{IH}^{-1} \tag{2}$$
$$\theta_1 = \frac{\theta\left((1-\delta_1)\gamma_{IR} + \delta_1\gamma_{IF}\right)}{\theta\left((1-\delta_1)\gamma_{IR} + \delta_1\gamma_{IF}\right) + (1-\theta)\gamma_{IH}} \tag{3}$$
$$\delta_1 = \frac{\delta\gamma_{IR}}{\delta\gamma_{IR} + (1-\delta)\gamma_{IF}} \tag{4}$$
$$\delta_2 = \frac{\delta\gamma_{HR}}{\delta\gamma_{HR} + (1-\delta)\gamma_{HF}}. \tag{5}$$

#### 2 Formulation of coupling parameters according to the generalised gravity model

We define $\sigma_{ij} \in [0,1]$ to be the proportion of epidemiologically relevant contacts that individuals from population $i$ have with individuals in population $j$, where $\sum_j \sigma_{ij} = 1$. We define the coupling according to the generalised gravity model, which we outline here.

We define $v_{ij}$ to be the the number of visits from population $i$ to $j$. According to the generalised gravity model this is proportional to $N_i^a N_j^b / d_{ij}^c$, where $d_{ij}$ is the distance between populations $i$ and $j$ and the parameters $a, b$ and $c$ are to be inferred. The coupling, $\sigma_{ij}$, should then be proportional to the fraction of individuals from population $i$ visiting population $j$, which is $v_{ij}/N_i$. In addition, we need to ensure that within population coupling $\sigma_{ii} = 1 - \sum_{j \neq i} \sigma_{ij}$ is always positive, placing a limit on the maximum size of external couplings. To this end, we normalise the proportion of visits by the maximum proportion over all populations; however, from this definition we have $\min_i \sigma_{ii} = 0$. Therefore, we introduce an additional scaling parameter $A \in [0, 1]$ such that $\min_i \sigma_{ii} \geq 0$. Combining each of these elements we define the coupling $\sigma_{ij}, i \neq j$, to be:

$$\sigma_{ij} = A \frac{N_i^{a-1} N_j^b / d_{ij}^c}{\max\limits_k \left( \sum\limits_{l \neq k} N_k^{a-1} N_l^b / d_{kl}^c \right)}, \tag{6}$$

where $a, b, c$ and $A$ are parameters to be inferred from the epidemiological dynamics.

To parameterise the spatial component of the model, we need an estimate of both population sizes and pairwise distances between populations. We use estimates of population sizes from global health bodies and define pairwise distances to be great-circle distances between populations (Table 1).

### 3 Prior distributions used in parameter inference

Parameters are inferred by calculating the weighted root mean square error between realised ($C^{sim}$) and observed ($C^{obs}$) cumulative confirmed cases in each of the sub-populations from $T_0 =$ until $T_1$ days later. Parameter inference is performed as a two step process, where in the first step parameter values are initially chosen from uniform prior distributions. We

| | | | Great-circle distance (km): | | | |
|---|---|---|---|---|---|---|
| Population | Pop. size | Location | Kinshasa | Wangata | Bikoro | Iboko |
| Kinshasa | $1.1 \times 10^7$ | 04°18′05″S 15°19′05″E | 0.0 | 583.4 | 520.9 | 530.2 |
| Wangata | $1.2 \times 10^6$ | 00°02′52″N 18°15′21″E | | 0.0 | 96.3 | 101.4 |
| Bikoro | 124,608 | 00°48′00″S 18°25′60″E | | | 0.0 | 17.5 |
| Iboko | 80,027 | 00°49′38″S 18°35′18″E | | | | 0.0 |

**Table 1.** A summary of the demographic data used to parameterise the spatial model. For each population of interest, we give approximate population size and location, from which we can derive pairwise great-circle distances between populations. Population size estimates are from census data, or, where not available, through other online sources.

use the following prior distributions for the

$$T_0 \sim \text{Uniform}\{4 \text{ April}, 11 \text{ May}\} \quad (7)$$

$$\hat{\beta}_0 \sim \text{Uniform}(0, 3) \quad (8)$$

$$T_C \sim \text{Uniform}\{11 \text{ May}, 11 \text{ May} + T_1\} \quad (9)$$

$$\delta \sim \text{Uniform}(0, 1) \quad (10)$$

$$a, b, A \sim \text{Uniform}(0, 1) \quad (11)$$

$$c \sim \text{Uniform}(0, 3). \quad (12)$$

### 4 Early case growth: additional results

Here we present additional figures for the simple model on $K = 3$ sub-populations (Iboko, Bikoro and Wangata) using data until 25 May 2018.

#### 4.1 Best-fit coupling estimates

We recombine the four spatial parameters ($a, b, c$ and $A$) to obtain meaningful distributions of the coupling between the sub-populations (Figure 1).

#### 4.2 Best-fit time series

We obtain time series fits to the observed cumulative cases and the final distribution of cases corresponding to 25 May 2018 for the metapopulation as a whole (Figure 2a) and for the four sub-populations separately (Figure 2b).

We compare the final distribution of the realised cumulative confirmed cases to the observed cumulative cases on 25 May 2018 (Figure 2, RHS). We estimate the total number of confirmed cases to be 37 (mean 37.0, 95% CI [30, 45]), compared to 35 observed confirmed cases. We can also look at the distribution of cases in each of the three sub-populations. In Wangata we estimate 5 confirmed cases (mean 4.9, 95% CI [3, 7]), compared to 4 observed cases; in Bikoro we estimate 14 confirmed cases (mean 13.6, 95% CI [9, 18]) compared to 10 observed cases; and in Iboko we estimate 18 confirmed cases (mean 18.5, 95% CI [14, 23]), compared to 23 observed cases. In all three sub-populations, the observed data fall within our 95% credible intervals.

### 5 Changes in transmission: additional results

Here we present additional figures for the breakpoint transmission model on $K = 3$ sub-populations (Iboko, Bikoro and Wangata).

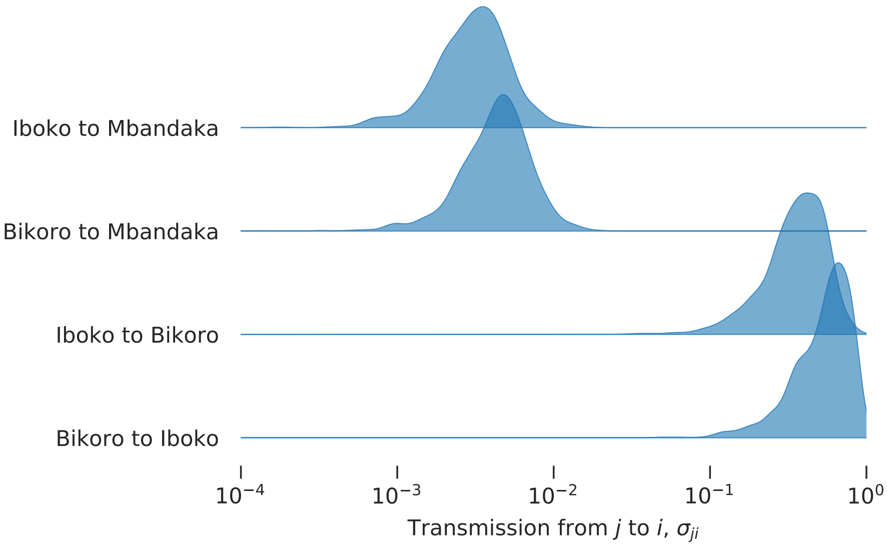


**Figure 1.** Select between-population coupling using the best-fit parameter estimates for $a, b, c$ and $A$. Here we refer to the transmission, which is the direction in which the disease moves; however the coupling is determined by the movement of susceptible individuals who bring infection back into their home population.

#### 5.1 Best-fit parameter estimates

We obtain best-fit estimates of the unknown parameters as additional confirmed cases are reported. Using data until 6 July 2018 we estimate the percentage reduction in transmission, $1-\delta$, to be 98.7% (95% CI [92.2%, 100%]) (Figure 3).

**(a)**

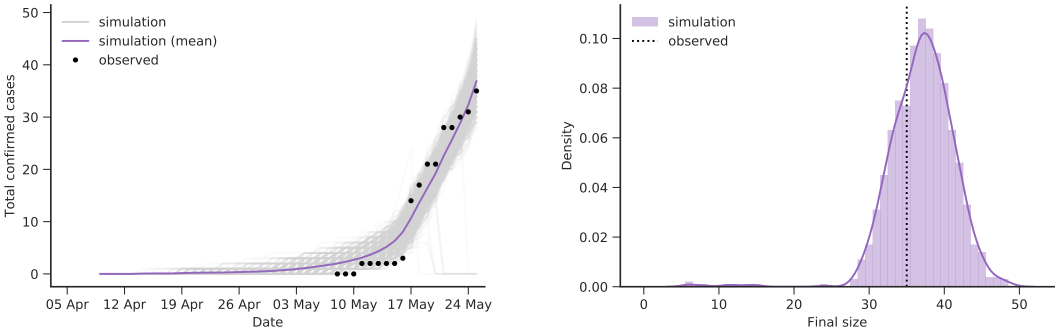


**(b)**

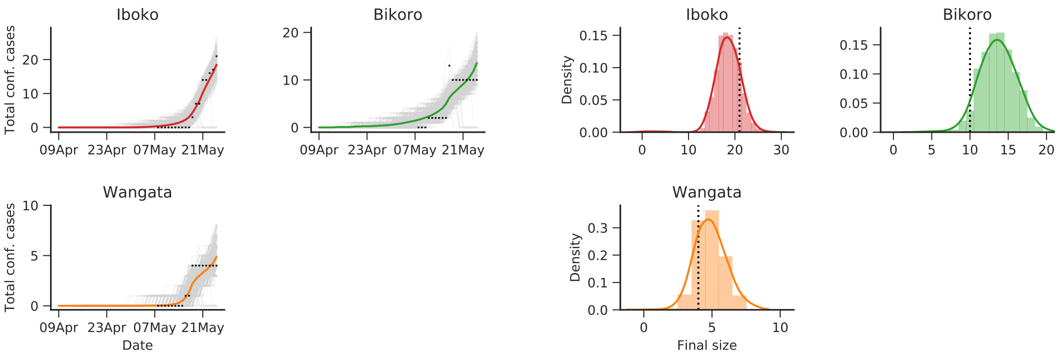


**Figure 2.** The epidemic curve and final distribution of cases for the 1,000 realisations with the smallest total error for the simple model on $K = 3$ sub-populations (Iboko, Bikoro and Wangata) up to 25 May 2018. Results are shown for (a) the metapopulation, summed over all sub-populations, and (b) the three sub-populations separately. On the left-hand side we show individual realisations of the outbreak (shown in light grey) plotting the cumulative number of cases moving from the infected to either the hospitalised or funeral class; the mean across all 1,000 realisations is plotted in colour and solid points denote actual cumulative confirmed cases. On the right-hand side we show the distribution of total cases from the 1,000 best realisations together with the reported value (shown as a dashed vertical line).

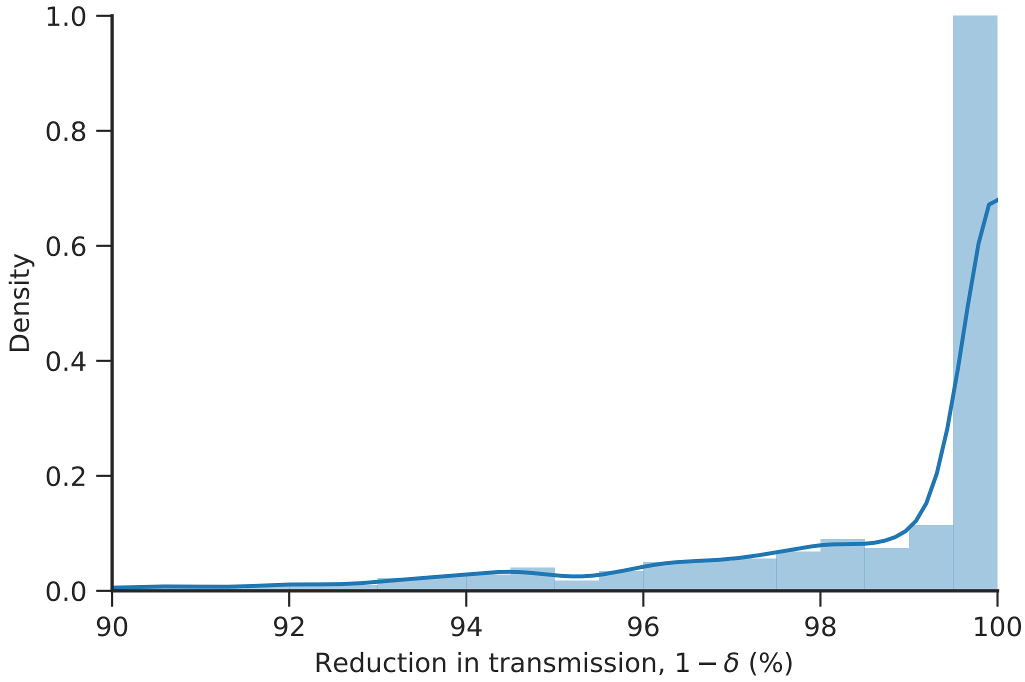


**Figure 3.** Posterior distribution for the percentage reduction in transmission $\delta$. These distributions are determined by the 1,000 best realisations fitted to data up 6 July 2018.
